# All for one or one for all? Disentangling the *Juncus bufonius* complex through morphometrics, cytometry and genomics

**DOI:** 10.64898/2026.02.24.707752

**Authors:** Billy Williams-Marland, Regina Berjano, Karin Tremetsberger, Jennifer Rowntree, Raúl Sánchez García, Casper H.A. van Leeuwen, Andy J. Green, Mª Ángeles Ortíz

## Abstract

*Juncus bufonius* L. *s.l.* is a species complex with several ploidy levels, for which species delimitation remains unclear due to a lack of reliable morphological characters and the paucity of molecular studies. To clarify taxonomic and geographic relationships in the complex, we combined genomic, cytometric and morphological data from a broad latitudinal range from England down to Spain. We collected morphometric and cytometric data from 31 populations, and genomic data were obtained through Hyb-Seq using the Angiosperm353 kit for a subset of individuals. These three datasets were combined to explore phylogenetic relationships, population structure, and the validity of four previously proposed morphospecies (*J. bufonius s.str.*, a hexaploid; *J. minutulus*, a tetraploid; and *J. ranarius* and *J. hybridus*, both diploids). Sequencing supported the separation of diploids and polyploids as two distinct taxa, but morphometric characters used previously to describe morphospecies showed continuous variation with no diagnostic value, and were not congruent with genomic and cytometric data. Polyploids likely originated through allopolyploidisation from diploids and tetraploids. Phylogenetic lineages were extensively mixed geographically, both for diploid and polyploid taxa, which suggests repeated long-distance dispersal events for both diploids and polyploids, and no separation of taxa by geography. Splitting of diploids into *J. ranarius* and *J. hybridus* was not supported. We recommend *J. ranarius* be treated as a synonym of *J. hybridus*, and that tetraploids and hexaploids be grouped under *J. bufonius*. The observed geographical patterns are consistent with high rates of seed dispersal by migratory waterbirds.

## INTRODUCTION

The *Juncaceae* Juss. are a cosmopolitan family of grass-like herbs that are commonly known as rushes. The genus *Juncus* L. is by far the largest and most widespread of the family with around 350 accepted species (World Flora Online, 2025) that are found in temperate to cold regions of the globe and more rarely in the tropics. While the genus is traditionally divided into two subgenera and 10 sections (Kirschner 2002a; b), molecular studies found this classification to be non-monophyletic (Drábková *et al*. 2003, 2004, 2006; Drábková and Vlček 2009; Drábková 2010). Efforts to regain monophyly have resulted in the proposal and creation of new genera such as *Oreojuncus* (Drábková and Kirschner 2013) or the more recent, but controversial, proposal to rearrange *Juncus* into six genera (Brožová *et al*. 2022; Proćków and Drábková 2023a; b; Elliott *et al*. 2023).

The *Juncus bufonius* L. polyploid complex (commonly known as toad rushes) is a taxonomically intricate group of species situated in *Juncus* sect. *Tenageia* Dumort. In the most recent revision of the family, Kirschner *et al*. (2002b) recognised six species in the complex: *J. bufonius* L. *s.str.*, *J. minutulus* (Albert & Jahand.) Prain, *J. ranarius* Songeon & E.P.Perrier, *J. hybridus* Brot., *J. sorrentinii* Parl. and *J. turkestanicus* V.I.Krecz. & Gontsch. Previous authors have also considered *J. foliosus* Parl. and *J. rechingeri* Snogerup to be closely allied to the complex (Cope and Stace 1978; Fernández-Carvajal 1982; Romero-Zarco 2010). However, the Plants of the World Online database (POWO 2025) lists 11 species, accepting three recent additions (Kirschner *et al*. 2004; Dobignard *et al*. 2010; Veldkamp 2014). Similarly, Proćków & Drábková (2023b) also accepted 11 species in their revision of the genus, although under new names. The centre of diversity of the complex lies within the West Mediterranean region (Kirschner 2002b). Nevertheless, several of these putative species are reported to have cosmopolitan or sub-cosmopolitan distributions, mainly, the two polyploids of the complex: *J. bufonius s.str.*, a hexaploid, and *J. minutulus*, a tetraploid. An important reason for this widespread distribution is that migratory waterbirds disperse large amounts of viable toad rush seeds through ingestion, transport and egestion in faeces (endozoochory) (Martín-Vélez *et al*. 2021; Navarro-Ramos *et al*. 2024; Jiménez-Martín *et al*. 2024). Toad rushes are typically found in open, wet habitats frequented by waterbirds that regularly move their seeds over distances of tens or even hundreds of kilometres (Martín-Vélez *et al*. 2021; Lovas-Kiss *et al*. 2023). Observation of such effective long-distance dispersal contrasts with the idea that the species complex consists of so many different species.

The delimitation of toad rush species is an enduring debate among ecologists and taxonomists. Despite extensive research (Cope and Stace 1978; Rooks *et al*. 2011), there remains a lack of clear and reliable diagnostic characters (Romero-Zarco 2008, 2012; Rudner and Deil 2011). Extreme morphological variability promotes disagreements among authors and contradictory results. In particular, Rooks et al. (2011) showed that the two polyploid cytotypes, were morphologically indistinguishable in central Europe, and argued that the tetraploid should be included within the taxonomic circumscription of *Juncus bufonius* instead of being segregated under the name of *J. minutulus*. Hence, the systematics of the complex remains unclear and controversial, especially due to the lack of molecular studies.

The systematics of polyploid complexes can be difficult to disentangle. Prezygotic and postzygotic barriers to reproduction often lead to speciation after whole genome duplication, *i.e.* polyploidy (Ramsey and Schemske 1998). However, these barriers can be permeable, allowing for interploidy reproduction, introgression and hybridisation which can lead to reticulate phylogenetic relationships that are challenging to resolve (Soltis *et al*. 2004; Parisod and Besnard 2007; Balao *et al*. 2016; Afonso *et al*. 2021; Karbstein *et al*. 2022; García-Cárdenas et al. 2025). Polyploidy can be divided into autopolyploidy (the duplication of a single species’ genome) or allopolyploidy (duplication following hybridisation). Both are considered to be important drivers of plant evolution, although the former has historically been overlooked (Spoelhof *et al*. 2017; Lv *et al*. 2024). Furthermore, polyploids have been regarded as evolutionary dead-ends with limited potential(Soltis *et al*. 2014). Recently formed polyploids, *i.e.* neopolyploids, face challenges such as genomic instabilities (Comai 2005; Soltis *et al*. 2014), higher extinction rates (Mayrose *et al*. 2011) or difficulties in establishment due to minority cytotype exclusion (Levin 1975). However, if they overcome these difficulties, polyploids possess extreme evolutionary potential (Soltis *et al*. 2014) often allowing them to inhabit novel niches and extreme environments (Comai 2005; Van De Peer *et al*. 2017, 2021; but see Mata *et al*. 2023). Moreover, polyploids can survive episodes of global climatic change or mass extinction, a further display of their adaptive potential (Van De Peer *et al*. 2021).

To date, most research on toad rushes has focused on diagnostic morphometric characters, leaving the systematics of the complex largely unresolved. Here, we aim to apply novel phylogenomic approaches to disentangle the relationships within the complex. Specifically, our objectives were to: (1) combine morphological, genomic and cytometric data to reconsider the validity of the proposed species of the complex; (2) compare genomic data over a broad latitudinal range in Europe to identify putative taxa and their distributions; (3) look for evidence that ploidy levels vary between habitats or latitudes; and (4) identify the origin and mechanism through which the polyploid taxa arose.

## MATERIAL AND METHODS

### Plant material

Specimens of *Juncus bufonius s.l.* were collected between 2021 and 2024 from margins of wetlands at 31 locations across Europe. Most locations hold high concentrations of migratory waterbirds, including species known to disperse *J. bufonius s.l.* seeds through endozoochory (Almeida et al. 2022). Locations were mainly in south-west Spain and England, with additional locations in Austria, Italy (Sicily), the Netherlands, Germany and the Czech Republic (Supporting Information Table S1).

Where possible, 10-20 mature individuals were collected in each location, and pressed or stored in bags with silica gel. For some individuals from each location, we attempted to identify morphospecies (see below) following keys in *Flora iberica* (Romero-Zarco 2010) and *Flora of the World* (Kirschner 2002b). Individuals with intermediate traits could not be assigned to a described morphospecies, and were classed as “uncertain”.

### Flow cytometry

Flow cytometry identified the ploidy level of some individuals from each population (Table S1). Cytometry samples were prepared with leaf tissue and the internal standard (*Solanum lycopersicum*; 2C = 2.59 pg). Measurements were taken on a CytoFLEX S flow cytometer with a 488 nm laser (Beckham Coulter, Fullerton, CA, USA). Total DNA content of each toad rush sample was calculated by multiplying the standard’s DNA content by the ratio between 2C peak positions of the sample and standard on the fluorescence intensity histogram. We tested the correlation between DNA content and latitude using Spearman’s correlation coefficient (ρ) and its significance with a linear model.

### Morphological study

The morphological study included 485 toad rush individuals from 17 sampled locations (Table S1) and 60 herbarium specimens identified as *J. minutulus* (Albert. & Jahand.) Prain (Supporting Information Table S2). The latter were included because few field specimens were assigned to that morphospecies.

Measurements were taken on mature individuals, i.e. those with dehisced capsules. The morphological variables included commonly used diagnostic characters (Kirschner 2002b; Romero-Zarco 2010) and were selected based on previous morphological studies (Van Loenhoud and Sterk 1976; Cope and Stace 1985; Rooks *et al*. 2011).

Quantitative variables included the length of the capsule (CL), the outer tepals (OTL) and the inner tepals (ITL). Additionally, ratios were calculated between CL and OTL (ROTC), CL and ITL (RITC), and OTL and ITL (ROTIT). Floral measurements were taken from approximately two flowers per individual (total of 1027 flowers). These were measured from photographs taken with a stereo microscope equipped with a HD camera (EZ4 W; Leica Microsystems) using Image J.JS software (Ouyang *et al*. 2019). CL, OTL and ITL were measured from the base to the tip of the structure, including the mucro or spike in the case of the capsule. Qualitative characters noted by visual inspection were seed coat morphology (smooth, striated or intermediate), and the disposition of the flowers in the inflorescence (solitary or clustered).

Data analysis was carried out in R v.4.4.2. (R Core Team 2024). We used statistical clustering and ordination techniques to see whether specimens segregated into morphological groups comparable to putative taxa within the complex. To avoid circularity, we did not assign these specimens to morphospecies prior to the analyses. Principal Components Analyses (PCA) were performed with the base *prcomp* function using scaled and centred data, retaining principal components that explained the most variance. Following this, to search for distinct groups we performed K-means clustering using the within cluster sum of squares method to determine the optimal number of clusters.

Additionally, we tested for morphological differences between the ploidy levels for a subset of individuals with morphometric and cytometric data. Univariate normality was violated according to Shapiro-Wilk tests. Kruskal-Wallis tests were used to test for significant differences between ploidy levels using the base stats package. Pairwise comparisons were performed using Dunn’s test from the FSA package (Ogle *et al*. 2023).

### Phylogenomics

The ingroup comprised 36 individuals from 19 sampled locations, representing all recorded ploidy levels (2x, 4x, 6x) (Supporting Information Table S4). The outgroup comprised four species from the cyperid clade: *Marsippospermum gracile* (Hook.f.) Buchenau and *Juncus gerardii* Loisel. from the *Juncaceae*, and *Carex multispiculata* Luceño & Martín-Bravo and *Hypolytrum nemorum* (Vahl) Spreng from the *Cyperaceae*. The two sedges were included to ensure proper rooting, given ongoing rearrangements of the *Juncaceae* genera. Outgroup sequences were acquired through the Tree of Life database (Baker *et al*. 2022) (Supporting Information Table S5).

Genomic DNA were isolated from leaf tissue using the DNeasy Plant Mini Kit (Qiagen, Hilden, NRW, Germany) following manufacturer’s instructions. Relative DNA concentration (ng/μL) was measured with a dsDNA Quantification Assay Kit using a QubitTM 3.0 fluorometer (Thermofisher scientific, Waltham, MA, USA). Library preparation and Hyb-Seq implementation were carried out following Pokorny *et al*. (2024).

Quality checks were performed on the demultiplexed paired-end reads with FastQC v.0.11.5 (Andrews 2010) and MultiQC v.1.14 (Ewels *et al*. 2016). FastP v.0.23.4 (Chen 2023) was then used to remove adapter sequences, poly-G and poly-X tails, and to trim low-quality bases (<Q20) in a 4-bp sliding window, after which sequences shorter than 40 bp were discarded. Sequence quality was then rechecked.

Nuclear and plastid genes and their flanking regions (*i.e.* supercontigs) were assembled and recovered with HybPiper v2.1.6 (Johnson *et al*. 2016). Filtered reads were mapped to nuclear and plastid targets with mega353.fasta (McLay *et al*. 2021) and a custom plastid target (Pokorny *et al*. 2024) using the BWA (Li and Durbin 2009) and DIAMOND (Buchfink *et al*. 2021) alignment algorithms, respectively. Summary statistics from HybPiper’s *stats* module were used along with the gene coverage score calculated with the max_overlap.R script from Shee et al. (2020) to assess the sequence assembly process for each gene and sample. Genes with a coverage score below ⅔ of the median were excluded from downstream processing, as were samples for which the number of recovered genes that were at least half as long as the target gene was below ⅓ of the median. Potentially paralogous gene sequences were detected with the HybPiper’s *paralog_retriever* module and were assessed when more than 2 sequence copies were recovered for a given gene. For this purpose, Maximum-Likelihood gene trees were constructed with IQ-TREE 2 (Minh *et al*. 2020) using the corresponding gene’s paralogs_all.fasta file. The trees were visualised with FigTree (Rambaut 2018), and, upon inspection, genes were excluded from the analyses when the paralogous sequences were located in distinct clades (Johnson *et al*. 2016; Johnson 2024). Finally, the nuclear and plastid supercontigs were retrieved using HybPiper’s *retrieve_sequences* module.

Nuclear supercontigs were aligned using MAFFT v.7.505 (Katoh and Standley 2013) with the *--auto* flag. Summary statistics were computed for the multiple sequence alignments (MSAs) with AMAS v.1.0 (Borowiec 2016) and were checked for alignment lengths and proportion of parsimony informative sites larger than the value of the Mdn+2SD, as well as genes with no informative sites. Following this, preliminary gene trees were inferred with FastTree 2 (Price *et al*. 2010) to detect and remove long branches with TreeShrink v.1.3.9 (Mai and Mirarab 2018), before refining the MSAs with trimAl v.1.4.1 (Capella-Gutiérrez *et al*. 2009) with the *-automated1* flag enabled. Summary statistics were then recalculated with AMAS. Final nuclear gene trees were inferred under Maximum-Likelihood with IQ-TREE 2 using ModelFinder (Kalyaanamoorthy *et al*. 2017) to select the optimal nucleotide substitution model. Branch support was tested with the non-parametric Shimodaira–Hasegawa-like approximate likelihood-ratio tests (SH-aLRT) with 1000 replicates. Following Simmons *et al*. (2023), unsupported branches in the concatenated gene trees were collapsed using Newick Utilities v1.6.0 (Junier and Zdobnov 2010). The output from this step was then used to infer the nuclear species tree under a multispecies coalescent model with ASTRAL-III (Zhang *et al*. 2018).

Plastid supercontigs were aligned with MAFFT, evaluated with statistics from AMAS, and MSAs were refined with trimAl following the same procedure as with the nuclear supercontigs. However, as the plastome represented a canonical coalescent gene (Doyle 2022), plastid supercontigs were concatenated and partitioned with AMAS before phylogenetic inference under a Maximum-Likelihood model with IQ-TREE 2. ModelFinder was used to select the optimal substitution model and branch support was tested with SH-aLRT and 1000 replicates.

### Analyses of Single Nucleotide Polymorphisms (SNPs)

Following the procedure described by Slimp *et al*. (2021) and Balant *et al*. (2024), a target file was created by extracting the longest sequence for each of the ingroup’s nuclear supercontigs. The paired-end filtered sequences, obtained during the phylogenomic analyses, were aligned to the target file with BWA and SAMtools v1.2.0 (Danecek *et al*. 2021) before performing the variant calls with GATK 4 (McKenna *et al*. 2010). Individual VCF files were combined to create the SNPs data matrix which was refined with BCFtools v1.18 (Danecek *et al*. 2021) with an initial hard filter (QD < 5.0 || FS > 60.0 || MQ < 40.0 || MQRankSum < -12.5 || ReadPosRankSum < -8.0 and *--missing-values-evaluate-as-failing*) and then with filters evaluating quality, missing data, sequencing depth and minor allele frequency, as well as removing multiallelic variants. Filtering steps were evaluated with summary statistics calculated with VCFtools v.0.1.16 (Danecek *et al*. 2011) and R (R Core Team 2024). The data matrix was also filtered for Linkage Disequilibrium using PLINK 2 (Chang *et al*. 2015) with the following parameters: *--indep_pairwise 50 5 0.2 --bad_ld*. The latter flag was enabled as the sample size was <50.

The filtered and unlinked SNPs were used to investigate the population structure. The Maximum-Likelihood approach implemented by ADMIXTURE (Alexander *et al*. 2009) was used to infer the samples’ ancestry proportions. These were obtained by first determining the optimal number of populations (*K*) through a 10-fold cross-validation process and selecting the values of *K* with the lowest and least dispersed errors. ADMIXTURE was then run independently for the selected values of *K* and the results were plotted in R. To explore the overall genetic variability of the population, a PCA was carried out with PLINK 2. Since PCA can fail to accurately describe differences between clusters in the population, a Discriminant Analysis of Principal Components (DAPC) was also performed (Jombart *et al*. 2010) with the adegenet R package (Jombart 2008)and was run independently for the optimal and suboptimal number of clusters as inferred by the *find.clusters* function from the package.

## RESULTS

### Flow cytometry

Total DNA contents followed a trimodal distribution with mean 2C peaks at 0.65 ± 0.04 pg, 1.1 ± 0.07 pg and 1.71 ± 0.11 pg (mean ± SD), matching previously documented values for the known ploidy levels in the complex. 2C values ranging between 0.55-0.79 pg were considered diploid; between 1.02-1.37 pg tetraploid; and between 1.49-2.08 pg hexaploid (Supporting Information Table S3; Fig. 1).

**Fig. 1.**
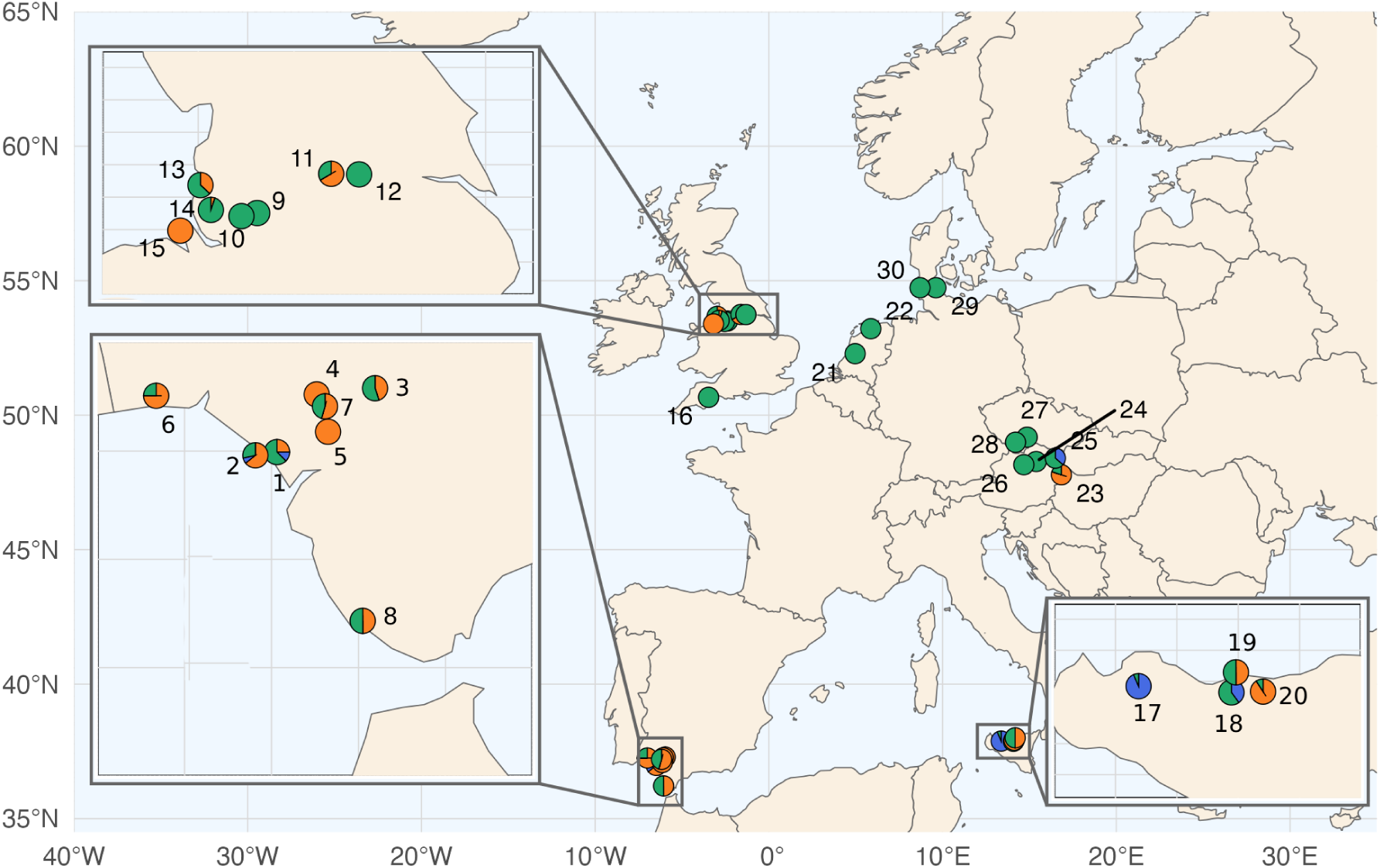
Distribution and proportion of diploids (orange), tetraploids (blue) and hexaploids (green) within sampled locations. Locations are listed as a number, see Supplementary Table S4 for location details and genome size, ploidy level and number of individuals in each of the cytometrically analysed populations.

Ploidy levels varied with latitude (ρ = 0.5459, p-value < 2.2e-16), with diploids increasing towards the south and hexaploids towards the north. Seven populations were purely hexaploid and only two purely diploid. Tetraploids were more frequent in Sicilian populations and none were solely comprised by this ploidy level. A majority of populations (14 of 23) showed mixed ploidy, mainly with diploids and hexaploids (Fig. 1).

### Morphology

Collected specimens were keyed out to four of the proposed morphospecies within the complex: *Juncus bufonius* L. *s.str.*, *J. minutulus* (Albert & Jahand.) Prain, *J. ranarius* Songeon & E.P.Perrier and *J. hybridus* Brot. However, many specimens (c. 13%) had intermediate characters, and could not be assigned to a single morphospecies with confidence.

The PCA was run including all quantitative variables (Fig. 2) in which the first and second principal components accounted for 43.2% and 34.8% of the variance, respectively. Within the PCA the individuals were distributed throughout the biplot in a sparse cloud surrounding the centre with no clearly delimited groups. Moreover, when coloured according to their assigned putative taxa, individuals were not separated into groups but were instead spread-out across the biplot. K-means clustering (using the two principal components retained from the PCA) identified the optimal and suboptimal number of clusters as 2 and 3 respectively (Supporting Information Figure S1). In both cases, K-means clustering was unsuccessful at grouping any of the putative morphospecies into a single cluster. Instead each cluster contained a mix of individuals assigned to different morphospecies (compare Fig. 2 and Fig. S1).

**Fig. 2.**
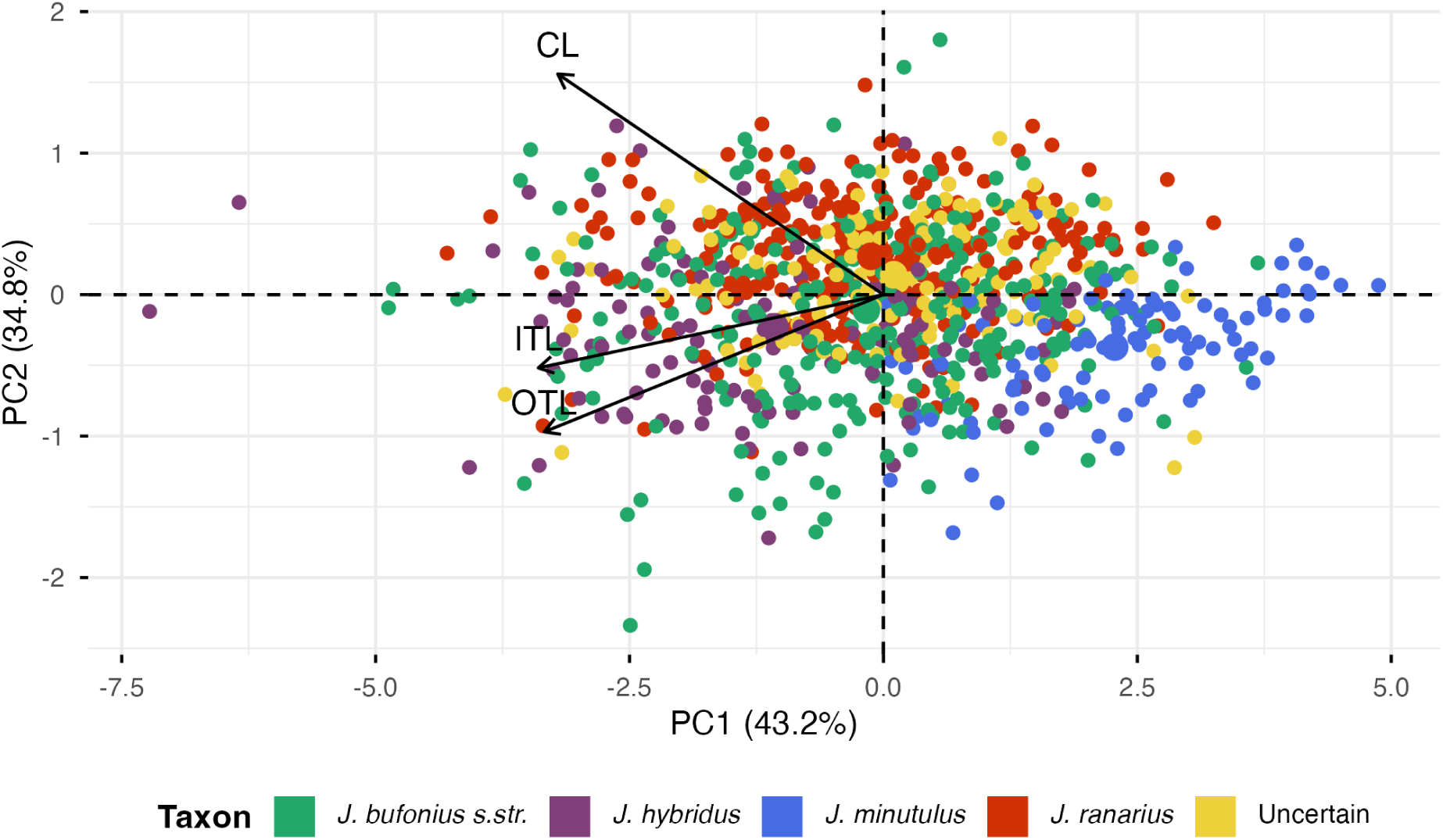
Biplot resulting from Principal Components Analysis using the capsule length (CL), inner tepal length (ITL) and outer tepal length (OTL) of 1028 flowers of *Juncus bufonius s.l.* Individuals are coloured according to their putative morphospecies identification.

When testing the morphology of the three ploidy levels, the ITL, RITC and ROTC reported significant differences according to Kruskal-Wallis tests (Supporting Information Table S6). Dunn’s tests revealed that all significant morphometric differences were between diploids and hexaploids, while tetraploids were statistically similar to both diploid and hexaploids (Supporting Information Table S7 and Fig. 3).

**Fig. 3.**
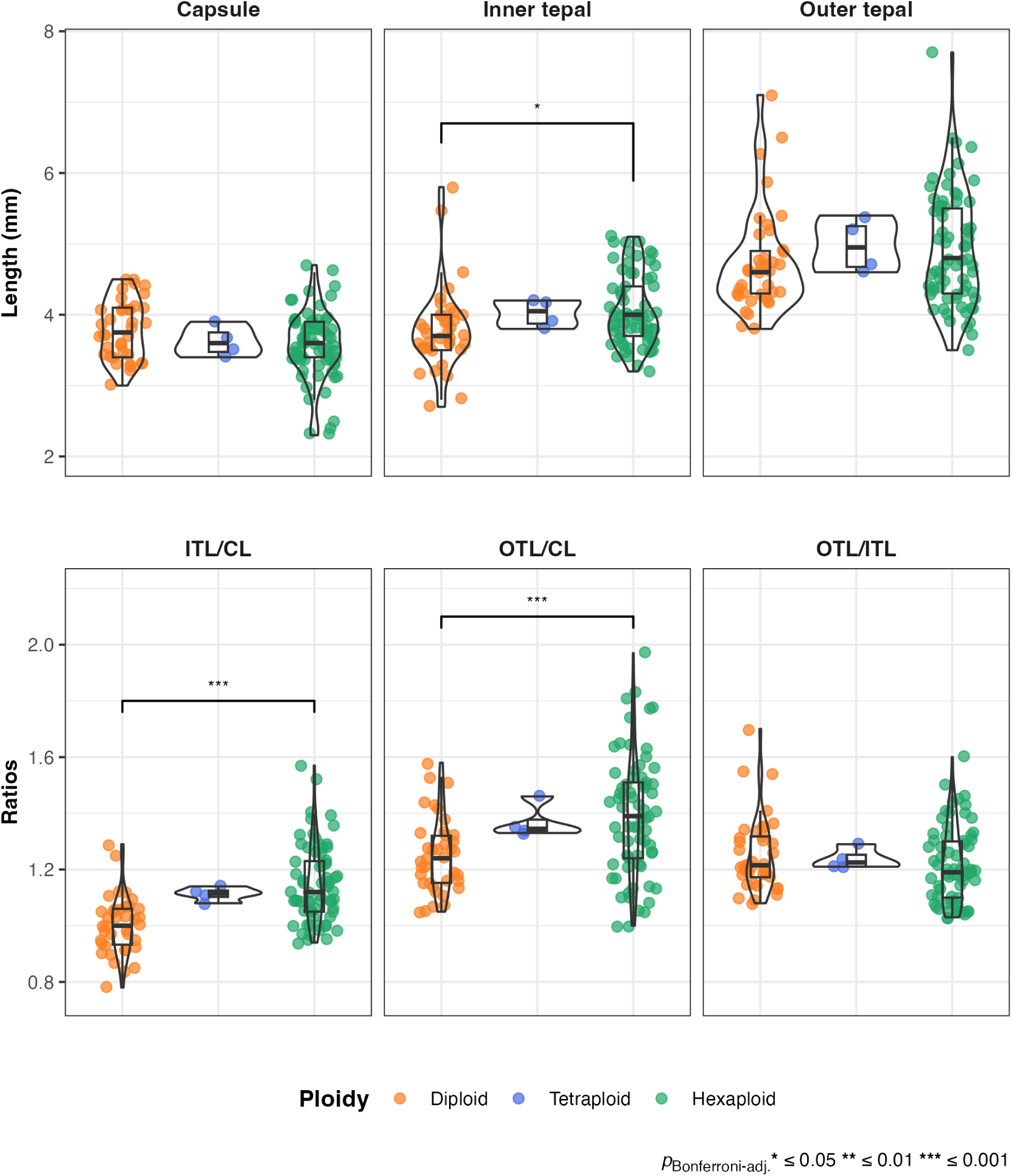
Violin and boxplots comparing morphological variables and ratios between individuals of *Juncus bufonius s.l.* of different ploidy levels. Individuals are coloured according to their ploidy level as determined by flow cytometry: diploids in orange, tetraploids in blue and hexaploids in green. Significant differences according to Dunn’s test are represented by an asterisk.

Regarding the qualitative characters, both seed ornamentation and disposition of the flowers were unreliable for separating putative species owing to overlap. Each of the four identified morphospecies possessed all three surface ornamentations (Supporting Information Table S8). Moreover, classifying the seeds into the three categories was complicated, as surface ornamentation presented in a continuum from smooth to striated. All morphospecies presented both solitary and clustered flowers (Supporting Information Table S9). Nonetheless, *Juncus bufonius s.str.* and *J. minutulus* presented mostly solitary flowers, while *J. hybridus* exhibited mostly clustered flowers. *J. ranarius* presented both solitary and clustered flowers in a similar proportion. In many cases, the distinction between solitary and clustered was not evident, especially in smaller specimens.

### Nuclear and plastid phylogenies

After trimming, an average of 5.8 million reads were obtained for each of the 36 individuals of the ingroup. HybPiper recovered, on average, 216 of the 353 possible nuclear genes and 64 of the 72 possible plastid genes per sample. After quality checks and filtering steps, one individual was excluded due to low gene recovery; 80 nuclear and 7 plastid genes were removed due to low coverage scores; and 8 plastid genes were removed due to the presence of paralogous sequences. In the end, 35 individuals remained in the ingroup; 272 nuclear supercontigs were used to infer the nuclear phylogeny, which yielded a species tree with a normalised quartet score of 0.56; and 57 concatenated plastid supercontigs were used to infer the plastid species tree.

The overall topology of the nuclear and plastid phylogenies was similar (Fig. 4) and both the outgroups and the ingroup were fully supported in both trees (LPP: 100 and SH-aLRT: 100). The two *Cyperaceae* formed a sister clade to the remaining taxa, all of which belong to the *Juncaceae* and clustered into their corresponding genera with *Juncus gerardii* as the sister taxon to the ingroup. The main dichotomy seen in both trees (Fig. 4) separated the ingroup into two principal clades, which were strongly but not perfectly related to ploidy levels. The diploid clade was fully supported in both trees (LPP:100 and SH-aLRT: 100), while the polyploid clade showed lower support in the nuclear tree (LPP: 50), but was fully supported in the plastid tree (SH-aLRT: 100).

**Fig. 4.**
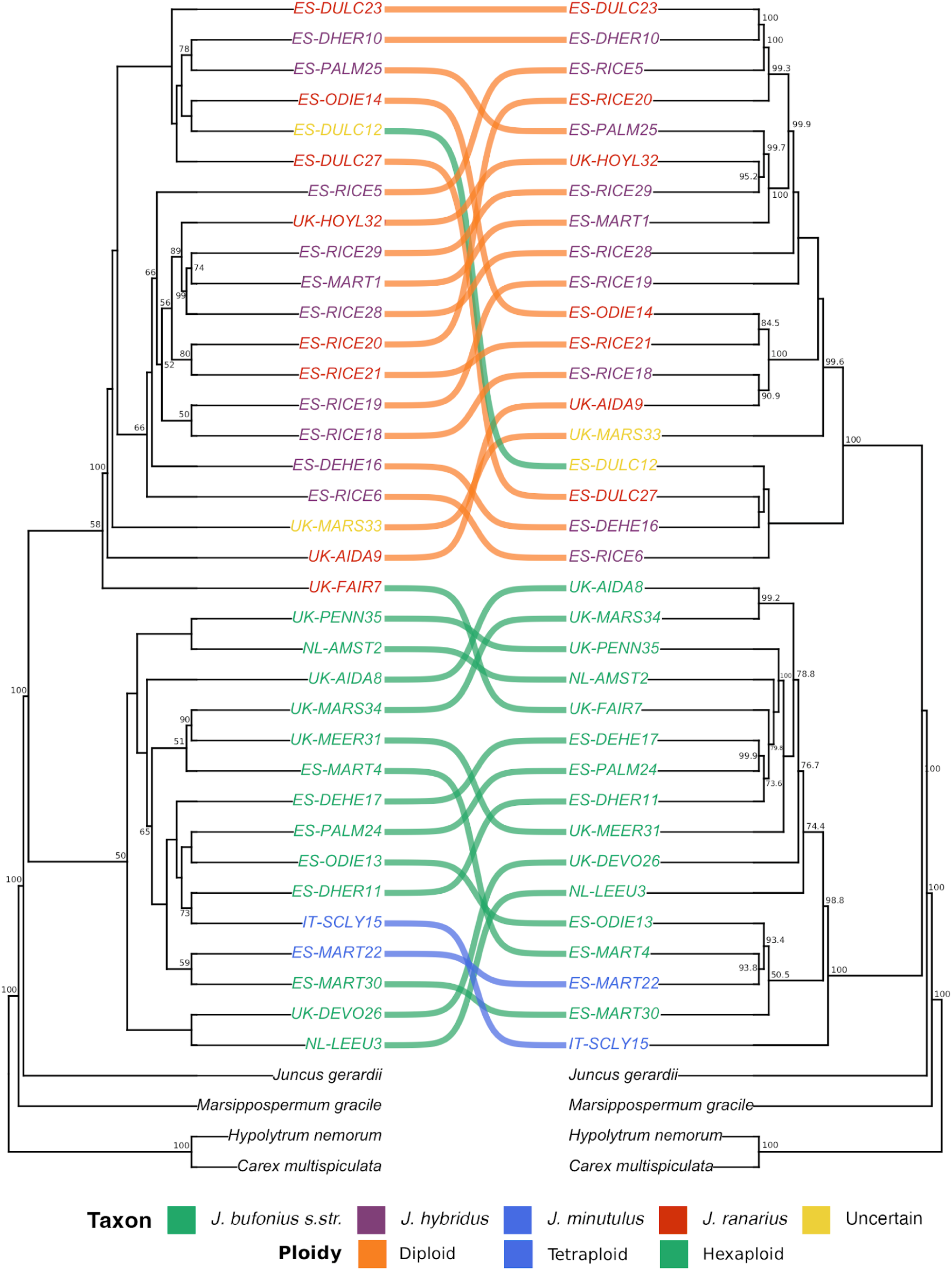
Cophylogenetic plot comparing the nuclear phylogeny (left) inferred under a multispecies coalescent model with ASTRAL-III from 272 Maximum-Likelihood nuclear gene trees and the plastid phylogeny (right) inferred from 57 concatenated plastid supercontigs under Maximum-Likelihood with IQ-TREE2. Branch support is measured in Local Posterior Probability (LPP) in the nuclear tree and in Shimodaira–Hasegawa-like approximate likelihood-ratio tests (SH-aLRT) in the plastid tree, and is shown when LPP and SH-aLRT ≥ 50. Individuals are coloured according to putative morphospecies identifications, and are named with the country code (ES: Spain, UK: The United Kingdom, NL: The Netherlands and IT: Italy), the sampled population (see Supporting Information Table S1 for population codes) and a number unique to each individual. Links are coloured according to the ploidy level of the individuals: diploids in orange, tetraploids in blue and hexaploids in green.

Despite overall similarities, the internal topology of the two main clades differed between nuclear and plastid phylogenies. Nevertheless, neither phylogeny grouped individuals into clades in a manner consistent with the assignment of morphospecies. In particular, in the diploid clades, *Juncus ranarius* and *J. hybridus* were consistently mixed together (Fig. 4).

#### The nuclear phylogeny

Individuals of the diploid clade were mostly distributed into two not well-supported subclades, except for two individuals from separate UK populations (UK-MARS and UK-AIDA), which occupied basal positions within the diploid clade. One of the subclades (LPP: 66) was mostly comprised of individuals collected from ricefields (ES-RICE), together with other individuals from nearby temporary marshes (ES-MART and ES-DEHE) and the Dee estuary in England (UK-HOYL). The remaining subclade (LPP < 50) was composed of individuals collected from other wetlands in SW Spain. Unexpectedly, a hexaploidy (ES-DULC12) was nested within the diploid clade and another (UK-FAIR7) occupied a sister position to the whole clade, although with low support (LPP: 58).

Within the polyploid clade, most hexaploid individuals clustered into one group that split into several subgroups of populations from different countries with varying support. The only two tetraploids were included in separate subclades of the polyploidy tree rather than grouping together (Fig. 4).

#### The plastid phylogeny

In the diploid clade within the plastid phylogeny, the individuals were mostly organised into several smaller highly supported (SH-aLRT > 99) subclades that showed a different pattern of phylogenetic relationships compared with the nuclear tree. However, the hexaploid ES-DULC12 was still situated within the diploid clade. In contrast, the hexaploid which had occupied a sister position to the diploid clade in the nuclear tree (UK-FAIR7) was now observed within the polyploid clade.

In the polyploid clade, the individuals were distributed between two subclades, except the tetraploid from Sicily (IT-SCLY15) which segregated from the clade into a highly supported sister position (SH-aLRT: 100). Although with low support (SH-aLRT: 50.5), one of the subclades was composed solely of individuals from SW Spain. The other subclade, exhibited higher support (SH-aLRT: 74.4) and contained individuals from the UK and the Netherlands with three individuals from SW Spain nested within (Fig. 4).

### Analyses of SNPs

In total, 9893 SNPs were recovered following variant calls for the 35 individuals of the ingroup. Following the filtering steps, 847 SNPs and 28 individuals remained in the population genomics analyses.

With ADMIXTURE, cross-validation revealed an optimal and suboptimal *K* value of 2 and 3, respectively (Fig. 5.A). For *K* = 2, all diploids, along with the aforementioned hexaploid, comprised one distinct genetic cluster. For *K* = 3, diploids split into two different groups, with several individuals exhibiting mixed ancestry. Most polyploids exhibited varying degrees of mixed ancestry between the inferred genetic groups for both values of *K*, except for one hexaploid (ES-DULC12) and one tetraploid (IT-SCLY15), which were assigned exclusively to one genetic group for both values of *K*.

**Fig. 5.**
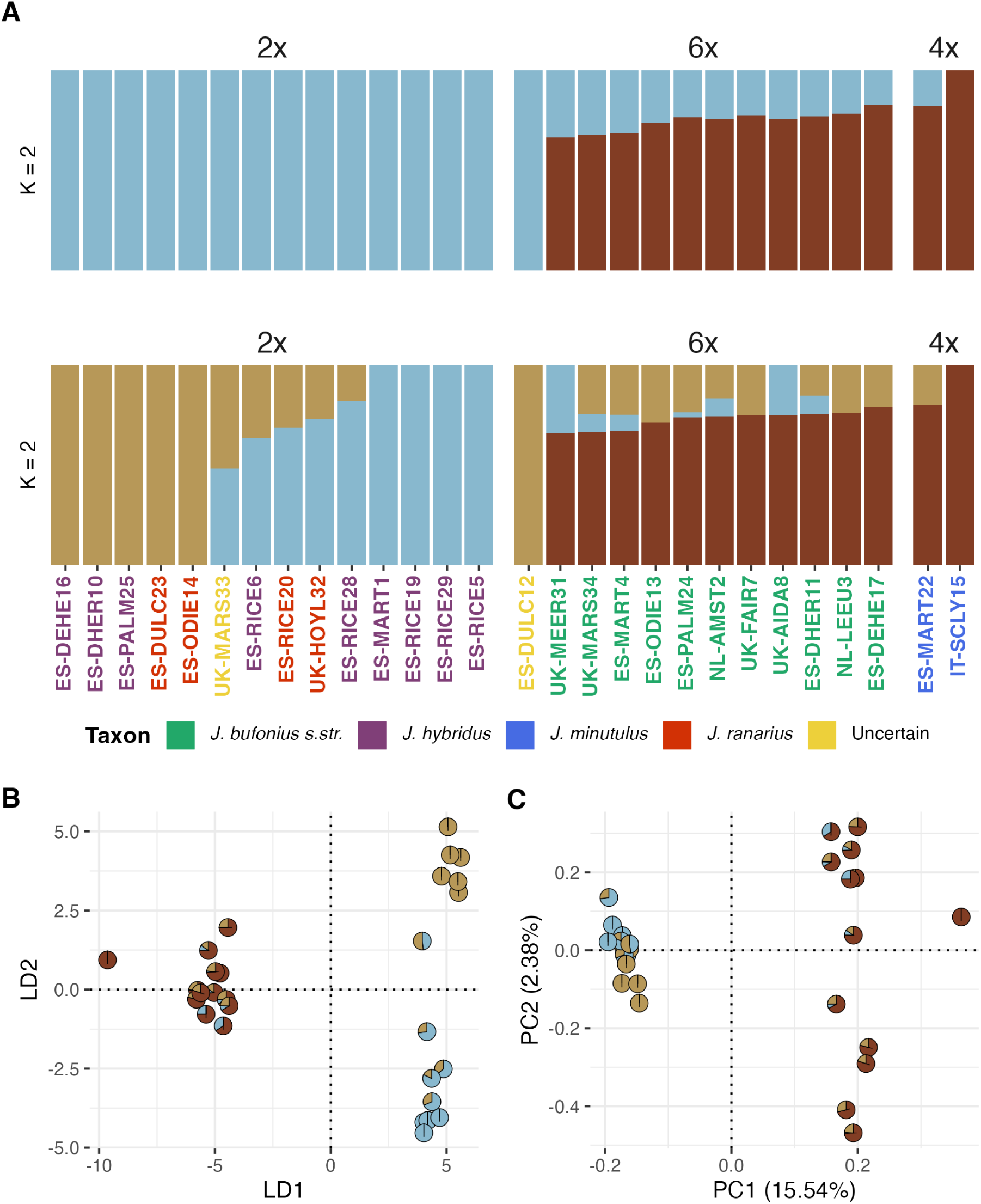
Results of the SNPs analyses carried out using 847 filtered and unlinked SNPs from 28 individuals of the ingroup used in the phylogenomics analyses. (A) Bar plots displaying the ancestry proportions of each individual inferred by ADMIXTURE for two and three genetic clusters, *K*. (B) Biplot of individuals using the first two dimensions of the Principal Components Analysis computed with PLINK2. (C) Scatter plot of individuals using the two linear discriminants obtained from the Discriminant Analysis of Principal Components for 3 genetic clusters. Individuals in Fig. B and C are represented by pie charts displaying their inferred ancestry proportions by ADMIXTURE for *K* = 3. Labels in Fig. A are coloured according to the putative morphospecies identification. See Supporting Information Figure S3 to see sample labels for Fig. B and C.

The K-means clustering algorithm used to identify groups for the DAPC also inferred two or three clusters within the population, the former having the lowest overall Bayesian information criterion (BIC = 77.69 for *K* = 2 and BIC = 78.29 for *K* = 3). The DAPC computed for *K* = 2 (Supporting Information Figure S2) produced the same pattern as LD1 in the DAPC for *K* = 3 (Fig. 5.C), which divided the individuals along the x axis in accordance to their ploidy level, except for the hexaploid ES-DULC12 which again grouped with the diploids. Furthermore, the diploids separated into two separate groups along the y axis (LD2), in a manner congruent with the ancestry proportion of each individual. An inverse pattern was also seen in the PCA (Fig. 5.B), in which the population once again separated into two groups according to ploidy level (PC1). However, in contrast to the DAPC, the polyploids were dispersed along the y axis (PC2) while the diploids comprised a relatively homogenous group. As before, this grouping is consistent with the genetic makeup of the individuals according to the ADMIXTURE results (*K* = 3; Fig. 5.A).

## DISCUSSION

To clarify the evolutionary relationships within the *Juncus bufonius* complex, this work integrates morphological, genomic and cytometric evidence. Our results suggest that the currently recognized morphospecies do not constitute coherent taxa, challenging the taxonomic treatment of the complex. Moreover, we found evidence of past polyploidisation and hybridisation events between ploidy levels, as well as multiple long-distance dispersal events for both diploids and polyploids. We also found evidence of habitat specialisation and possible local adaptation among diploids.

### Phylogenetic and taxonomic relations

This study is the first to apply genomic tools to the *Juncus bufonius* complex, in order to clarify relationships among putative taxa of the complex. The nuclear and plastid phylogenies (Fig. 4) were largely congruent with each other and revealed two independent, well-supported clades: diploids and polyploids. Likewise, the analyses of SNP data through ADMIXTURE, PCA and DAPC (Fig. 5) also discerned two principal genetic clusters within the population that related to ploidy level. Additionally, the separation of diploids and polyploids, more specifically the hexaploids, appeared to be supported by some differences in average values for morphological characters (Fig. 3), although extensive overlap in measurements remained between the ploidy levels. Collectively, these results support the separation of the diploids and polyploids as different taxa, but they do not support further splitting of taxa within the polyploid and the diploid levels. Many previous authors have included the diploid species within the taxonomic circumscription of *J. bufonius* as infraspecific taxa (*e.g.* Buchenau 1906; Husnot 1908; Maire 1957; Blanca López *et al*. 2009).

Toad rushes showed high morphological plasticity. We could not distinguish any morphogroup which correlated with previously described species. Morphology was extremely variable with no objective manner of grouping individuals into morphotypes. There were no appreciable discontinuities in the distribution of the variables’ values that support the division into multiple taxa (Fig. S4). The general aspect and measurements of structures even varied greatly within the same individual and, in many cases, could be assigned to different morphospecies based on the literature.

Hence, we found no objective support for length measurements, ratios and qualitative characters used to distinguish the species of the complex (Kirschner 2002b; Romero-Zarco 2010). Previous studies lack consensus regarding diagnostic characters. van Loenhoud & Sterk (1976) and Mičieta & Mucina (1983) reported the ability to distinguish diploids, tetraploids and hexaploids based on morphological features including the length of the capsule and the inner tepals. Similarly, Holub (1976) argued that *Juncus minutulus* can be separated from *J. bufonius s.str.* and *J. ranarius* bythe smaller size of some reproductive structures. In contrast, Cope & Stace (1978, 1983) included *J. minutulus* within the circumscription of *J. bufonius* as they were unable to separate them by morphological traits. Likewise, Rooks et al. (2011) were unable to separate cytometrically determined tetraploids and hexaploids by morphology, and found no support for *J. minutulus*.

As for the apparent presence of *Juncus ranarius* and *J. hybridus* within our samples, according to the character values from the classification keys, the phylogenomic and morphological data do not support their existence. The plastid phylogeny (Fig. 4), PCA (Fig. 5.B) and ancestry proportions (K = 2; Fig. 5.A) depicted a relatively homogenous diploid group with no support for the existence of two diploid taxa in the sample. Diploid individuals classified as *J. ranarius* and *J. hybridus* were mixed together within subclades of the plastid tree (Fig. 4). Morphological data provided no support for this splitting into two morphospecies (Fig. S1) and specimens assigned to either one overlapped within the space of the PCA (Fig. 2). Previous authors commented on such a morphological similarity and the difficulty in distinguishing one from another (Segal 1960; Romero-Zarco 2010). Altogether, our study points towards the existence of a single diploid taxon within our sample. We propose that *J. ranarius* be considered a synonym of *J. hybridus*, the latter being the older name. However, given the presence of other diploid taxa in the complex, a revision of the current taxonomic treatment of toad rushes may be warranted. Further research is desirable to collect genomic and other data from a larger sample size from a wider geographic distribution, including those areas where other species of the complex have been described, *e.g. J. maroccanus* in Morocco, *J. batrachium* in Borneo or *J. rechingeri, J. foliosus* and *J. sorretinii* in the Mediterranean Basin (Kirschner *et al*. 2002b; Kirschner *et al*. 2004; Veldkamp 2014).

In relation to the polyploid group, we found evidence to support a hybrid origin for most of the hexaploids, as the ancestry proportions inferred by ADMIXTURE (Fig. 5.A) showed the genetic makeup of these to be a mix of the diploid and tetraploid clusters. Cope & Stace (1985) were the first to propose that *Juncus bufonius s.str.*, the hexaploid, was an allopolyploid that had originated through the backcrossing of tetraploids with their diploid progenitors. Furthermore, the higher proportion of shared ancestry between hexaploids and tetraploids within ADMIXTURE may be indicative of the former backcrossing with tetraploid parents. Further insight into the emergence of the allohexaploids can be gained from the plastid phylogeny (Fig. 4). The appearance of two hexaploid subclades within the tree matching geographic regions may be indicative of different lineages. Polytopic and recurrent polyploid formation is a common occurrence in other complexes, and has the potential to increase genetic variability of polyploids, with important consequences such as variability in floral morphology (Ownbey 1950) or wider ecological breadth (Meimberg *et al*. 2009; Cuadrado *et al*. 2017). Moreover, polyploids of different origins can be interfertile (Doyle *et al*. 1999; Modliszewski and Willis 2012), which can aid in the propagation of variability through the species. Interbreeding polyploids may have arisen multiple times from different diploid progenitors. However, the present study did not include all the diploids of the complex (see above).

Although the majority of polyploids appear to be hybrid in origin, we could not rule out other possible mechanisms or origin, such as autopolyploidy. This was particularly evident in the case of the hexaploid ES-DULC12 that consistently clustered with the diploid individuals in both nuclear and plastid phylogenies (Fig. 4), in ADMIXTURE (Fig. 5.A), and also in the PCA and DAPC (Fig. 5.B and C). Autopolyploids within the toad rush complex have the potential to complicate the identification of morphospecies as can be morphologically similar to their diploid progenitors (Soltis *et al*. 2007; Barker *et al*. 2016; Spoelhof *et al*. 2017).

The origin of the tetraploids remains unclear. Like the allohexaploids, the Spanish tetraploid (ES-MART22) showed mixed ancestry inferred by ADMIXTURE (Fig. 5). On the other hand, the tetraploid IT-SCLY15, appeared to be genetically distinct from all other individuals. In the plastid phylogeny (Fig. 4) it adopted a sister position to the remaining polyploids and was assigned exclusively to a distinct genetic cluster by ADMIXTURE (Fig. 5.A). This could indicate that it originated from the other diploids of the complex, either through auto- or allopolyploidy. However, a more extensive sampling of both diploid and tetraploid lineages would be required to confirm the parental contributors to the tetraploids.

### Geographic relationships

The nuclear and plastid phylogenetic trees (Fig. 4) reflect high connectivity between the sampled locations, notably between Spain, the UK, and the Netherlands. Within the plastid phylogeny, subclades generally comprised individuals from all three countries and, on occasion, highly-supported sister sequences belonged to different countries (*e.g.*, the diploids UK-HOYL32 and ES-RICE29 with a SH-aLRT of 95.2). The relationship between these locations is likely driven by long-distance dispersal via migratory waterbirds. Such zoochorous seed dispersal provides the greatest dispersal distances in comparison to other vectors, such as anemochory or hydrochory (Navarro-Ramos *et al*. 2022; Jiménez-Martín *et al*. 2024). Indeed, the maximum dispersal distance by wind has been estimated at only 100m, compared to >100km by birds, even outside the migratory period (Martín-Vélez *et al*. 2021).

Long-distance dispersal may also explain, at least to some extent, the observed cytogeographical pattern in the sampled locations, specifically, the increase in hexaploids towards northern latitudes (Fig. 1). The more plausible scenario could be the dispersal from the Mediterranean region —the complex’s centre of diversity— to new environments, which may have allowed the polyploids to colonise novel niches that their diploid progenitors could not (Mummenhoff and Franzke 2007). Long-distance dispersal is considered more likely to succeed in polyploids than in diploids (Linder and Barker 2014). In our case, there is no good reason to expect greater dispersal limitation for diploids than for polyploids, since both frequent habitats where their seeds are ingested and dispersed by waterbirds. Indeed, some diploid plants found in the UK are closely related to others from Spain. Nonetheless, other factors such as climate may influence the establishment of the diploids in northern locations, as polyploids can be more cold tolerant (Van De Peer *et al*. 2021).

The establishment and persistence of polyploids depends on the interplay of several key factors (Fowler and Levin 2016). In stable environments, diploids generally outcompete their polyploid progeny on account of local adaptations, favouring niche differentiation between the two (Blaine Marchant *et al*. 2016; López-Jurado *et al*. 2019). In toad rushes, Rooks et al. (2011) considered the ecological preferences of the two polyploids to be identical. In contrast, in the Netherlands, van Loenhoud & Sterk (1976) reported tetraploids to prefer undisturbed environments, and hexaploids to prefer strongly disturbed habitats. We found different cytotypes present in many of our sampling localities (Table S3). The polyploids are wholly sympatric with the diploids of the complex, while also being the most widely distributed, as we also observed (Fig. 1). That being said, we also found purely diploid populations in the ricefields of SW of Spain plus a coastal population in the NW of the UK (Fig. 1). Furthermore, the ricefield diploids were genetically distinct from other diploids according to the genomic analyses (for *K* = 3, Fig. 5) and constituted a separate subclade within the nuclear phylogeny (Fig. 4). The sampled ricefields are surrounded by natural wetlands (Green *et al*. 2018) containing mixed ploidy populations, and shared waterbird populations dispersing seeds between the two types of environments (Jiménez-Martín *et al*. 2024). Hence, diploids likely outcompete polyploids in ricefields, owing to higher competitiveness under more stable conditions. The sampled ricefields have a relatively stable hydroperiod and are inundated annually for rice culture between May and October (van Rees et al. 2021). Toad rushes exploit this moist soil habitat during winter and spring months between autumn harvest and subsequent rice sowing. In contrast, surrounding populations in natural marshes depend directly on rainfall fluctuations, with strong and unpredictable variation in hydroperiod and salinity (Green *et al*. 2024). Under these circumstances, polyploidy can be advantageous. For example, in the *Dianthus broteri* complex, individuals of lower ploidy levels were more competitive under high water availability, while the opposite was true under low water availability (Rodríguez-Parra *et al*. 2024).

## CONCLUSIONS

This study provides valuable insight into the evolutionary history of the *Juncus bufonius* polyploid complex and highlights incongruences of its current taxonomic treatment. Genomic data supported the separation of diploids and polyploids as distinct taxa, but not any further taxonomic separations within each cytotype. Morphological data did not support splitting the specimens into the four currently recognised morphospecies. These findings support the grouping of hexaploids and tetraploids under the same taxon, *i.e. J. bufonius s.str.,* as proposed by Rooks *et al*. (2011). Similarly, the presence of two diploid taxa (*J. ranarius* and *J. hybridus*) was also not supported and we recommend *J. ranarius* be considered a synonym of *J. hybridus*. Although, further phylogenomic studies are required before full taxonomic revision of the complex. Future work should endeavour to include a larger sample size, as well as the remaining diploid taxa, from a wider range of locations throughout their distributions. Most polyploids seemingly originated through the hybridisation of the diploid and tetraploid gene pools. However, our results point to a possible origin in other diploids not covered by our sampling efforts. Moreover, the clustering of a hexaploid with the diploids suggests polyphyly and other origins for the polyploids of the complex. Finally, evidence for ecological differences were found between ploidy levels, specifically, adaptation among diploids to more predictable environments provided by ricefields. The *J. bufonius* complex could constitute an interesting system for further studies of polyploid dynamics and competition.

## DATA AVAILABILITY

The data underlying this article (morphometric, cytometric and, nuclear and plastid filtered reads, captured sequences, intermediary files and summary statistics for the phylogenomic and SNPs analyses) are available from Zenodo at [link]. The scripts used for the phylogenomic analyses are available from the GitHub repository at [link].

## SUPPORTING INFORMATION

Fig. S1. K-means clusters for K = 2 and 3 plotted on PCA biplot.

Fig. S2. Discriminant Analysis of Principal Components computed for K = 2 density plot.

Fig. S3. (A) Discriminant Analysis of Principal Components and (B) Principal Components Analysis from Fig. 5 A and B with sample labels.

Fig. S4. Violin plots of quantitative morphological variables and ratios.

Table S1. Population codes, habitat coordinates, voucher information and sample size in analyses for sampled locations.

Table S2. Herbarium specimens of *Juncus minutulus* (Albert & Jahand.) Prain used in the morphological study.

Table S3. Average genome size and number of individuals (*n*) per ploidy level in populations sampled for flow cytometry.

Table S4. Putative morphospecies identification, DNA content and ploidy level of the ingroup individuals of the phylogenomic analyses.

Table S5. Outgroup sequence information. Table S6. Results from Kruskal-Wallis tests.

Table S7. Results from Dunn’s pairwise comparisons between ploidy levels.

## AUTHOR CONTRIBUTIONS

B.W-M.: Fieldwork, data analysis and lead author. R.B.: Hyb-Seq implementation, project support and editing. K.T.: Flow cytometry, fieldwork, project support and editing. J.R.: Project support, editing and fieldwork. R.S.G.: Flow cytometry, editing and fieldwork. C.H.A.v.L.: Project support, editing and fieldwork. A.J.G.: Project conception and support, funding acquisition, fieldwork and editing. M.A.O.: Hyb-Seq implementation, project conception and support, fieldwork, editing and supervision.

## ACKNOWLEDGEMENTS

Sampling permits were provided by the Junta de Andalucía (project 2021/13), English Nature, The Wildfowl & Wetlands Trust, and The Royal Society for the Protection of Birds, and all necessary sampling permissions were obtained prior to sampling elsewhere. Genomic libraries were prepared at the laboratory facilities of the Herbarium and the Biology Research Services (CITIUS, University of Seville). Flow cytometry was performed at the Universität für Bodenkultur Wien (Austria). Bioinformatic workflows were implemented in the Hercules High Performance Computing cluster at the Centro Informático Científico de Andalucía (CICA). This work was part of a MSc degree (Máster Universitario en Biología Avanzada: Investigación y Aplicación, Universidad de Sevilla).

The authors would like to thank Iciar Jimenez Martín for her help with the plant sampling, Lisa Pokorny for her help with the Hyb-Seq workflow, Francisco García Cárdenas for his help with the SNPs workflow, and David Doblas Pruvost and Rafael González Albadalejo for their help and advice.

