## Supporting information for "All for one or one for all? Disentangling the *Juncus bufonius* complex through morphometrics, cytometry and genomics"

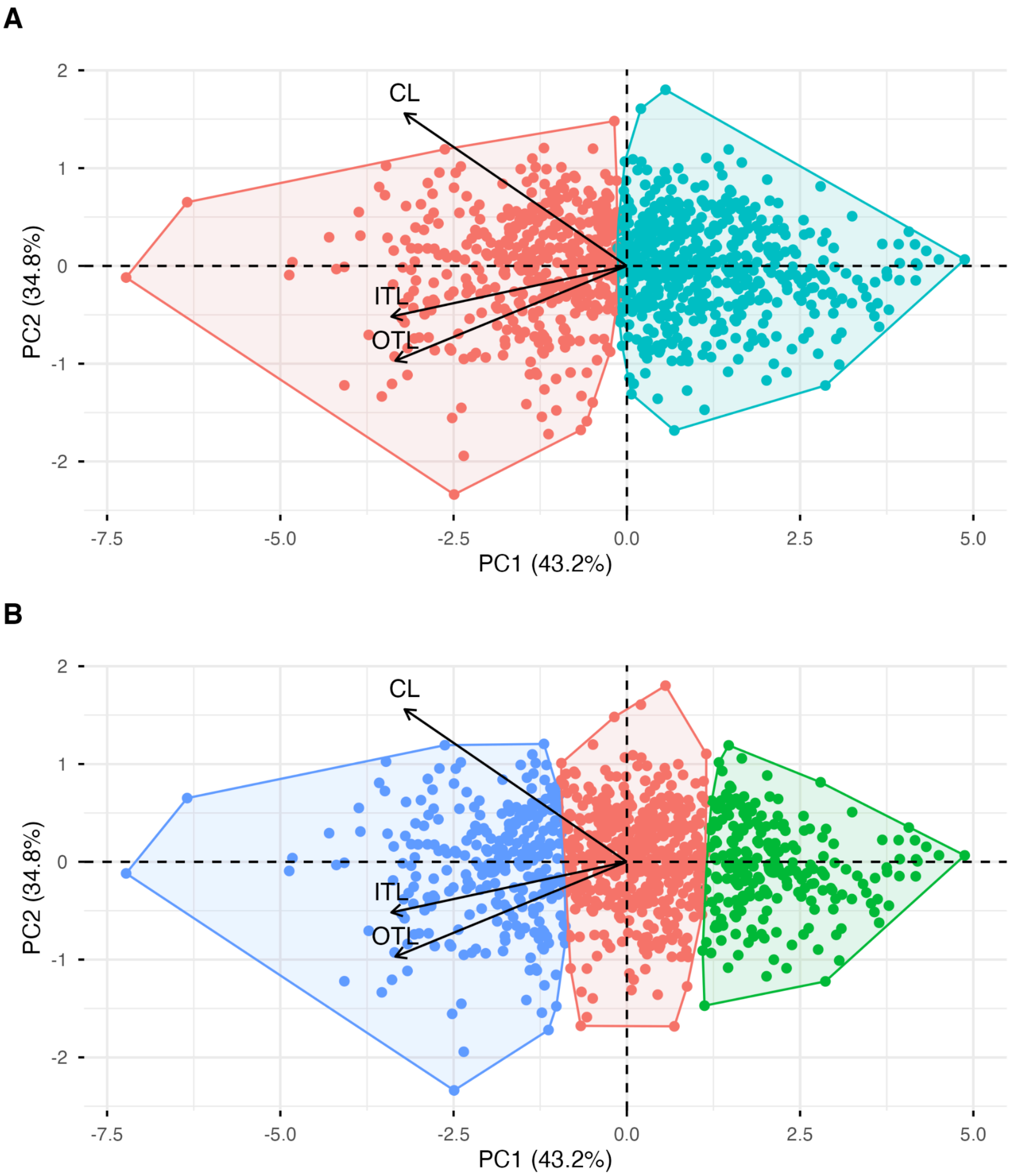


**Fig. S1.** PCA biplot from figure S2 with plotted K-means clusters for (A) $K = 2$ and (B) $K = 3$.

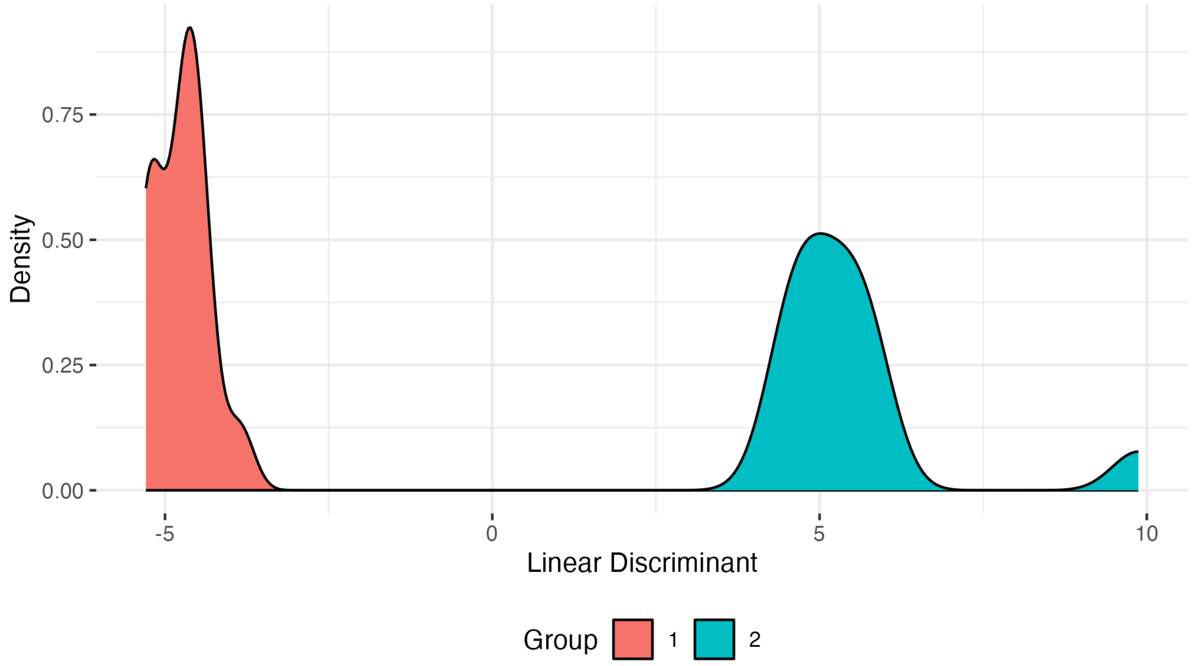


**Fig. S2.** Density plot of individuals according to the linear discriminant (LD1) obtained from the Discriminant Analysis of Principal Components for 2 genetic clusters using 847 filtered and unlinked SNPs from 28 individuals of the ingroup used in the phylogenomics analyses.

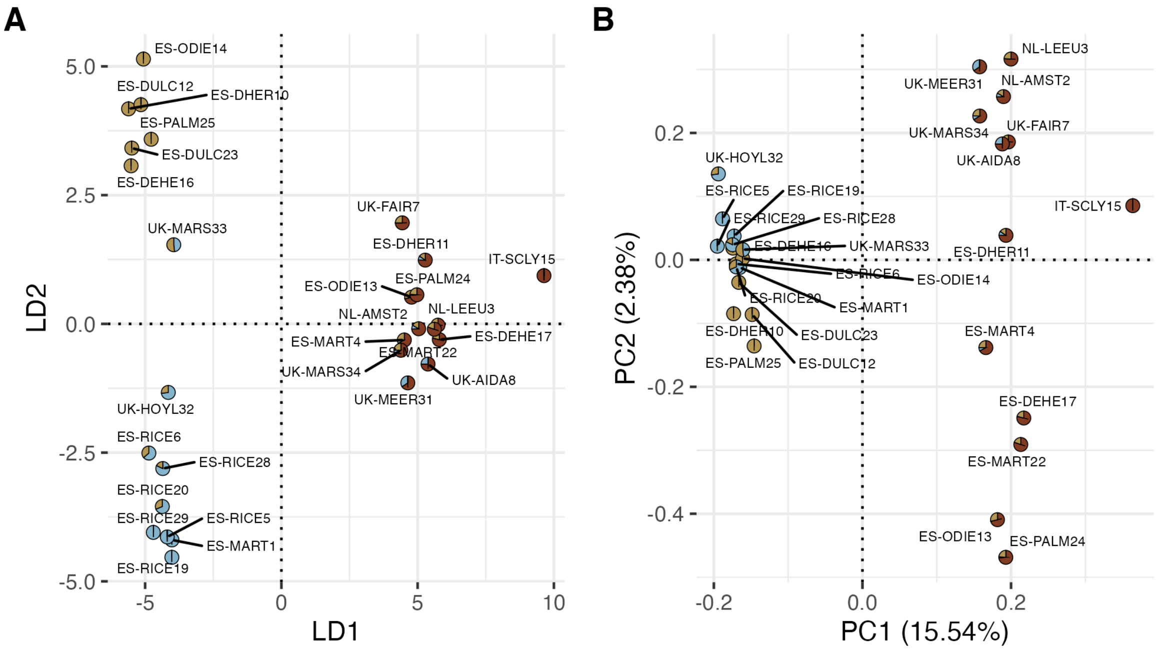


**Fig. S3.** (A) Discriminant Analysis of Principal Components and (B) Principal Components Analysis from Fig. 5 A and B with sample labels.

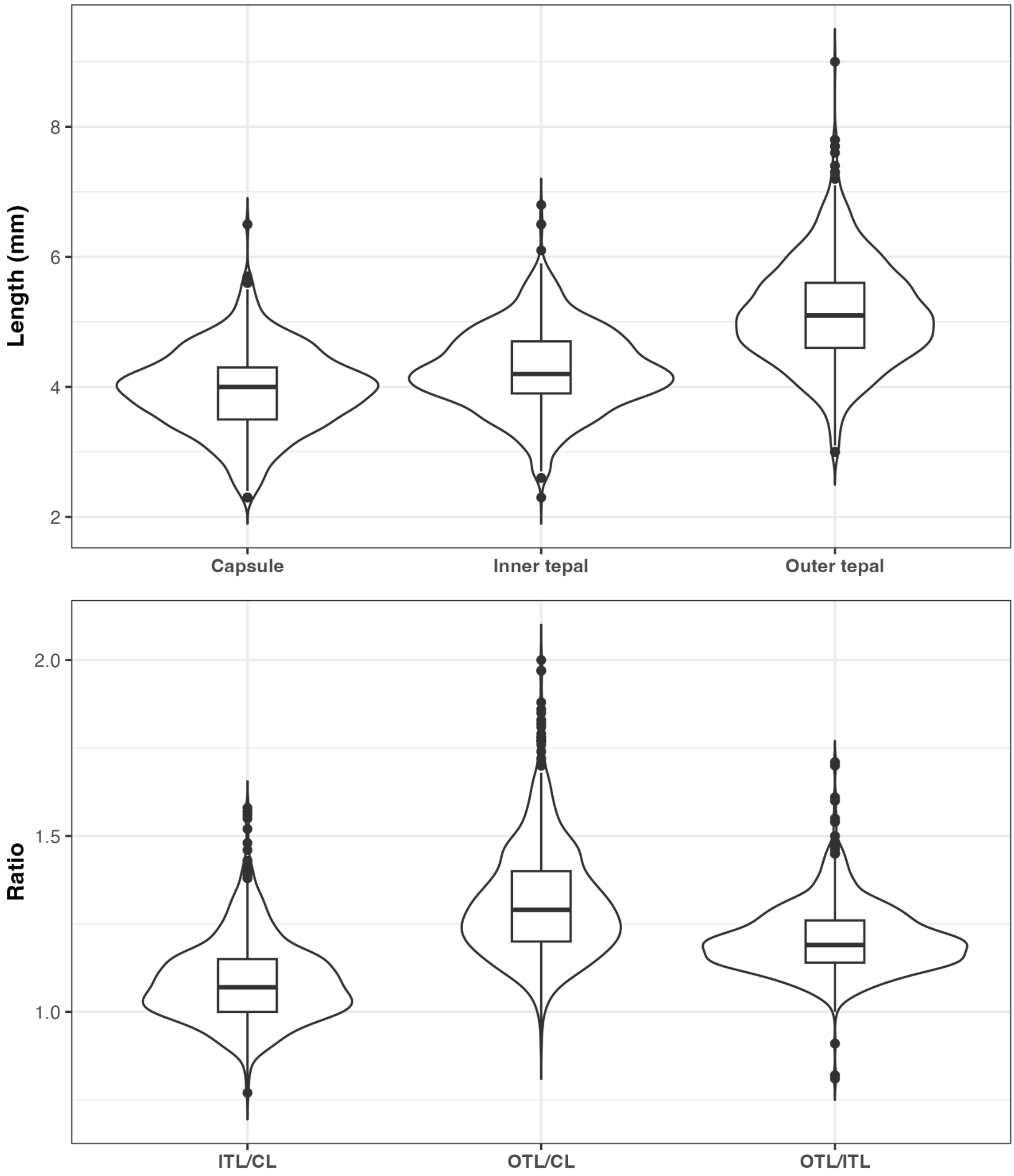


**Fig. S4.** Violin plots of quantitative morphological variables in mm and calculated ratios.

| **Supplementary Table S1.** Population codes, habitat, coordinates and sample size (*n*) in analyses for sampled locations. | | | | | | | |
|---|---|---|---|---|---|---|---|
| **Population code** | **Locality** | **Habitat** | **Coordinates** | **Herbarium voucher** | ***n* cytometry** | ***n* morphology** | ***n* phylogeny** |
| 1 ES-MART | Spain. Huelva, Almonte. Caño del Martinazo, Parque Nacional de Doñana. | Seasonal marsh, natural. | 36.99500, -6.45666 | SEV | 16 | 30 | 4 |
| 2 ES-DULC | Spain. Huelva, Almonte. Laguna Dulce, Parque Nacional de Doñana. | Semi-permanant dune pond. | 36.98109, -6.48487 | SEV | 14 | 34 | 3 |
| 3 ES-DHER | Spain. Seville, Dos Hermanas. Canal del Bajo Guadalquivir. | Temporary lagoon. | 37.29103, -5.96936 | SEV | 20 | 63 | 2 |
| 4 ES-RICE | Spain. Seville, La Puebla del Río. | Ricefields between harvest and sowing. | 37.26097, -6.07966 | SEV | 39 | 505 | 4 |
| 5 ES-RICE | Spain. Seville, Isla Mayor. | | 37.09083, -6.13778 | SEV | 36 | 145 | 4 |
| 6 ES-ODIE | Spain. Huelva, Aljaraque. Paraje Natural Marismas del Odiel. | Salt marshes. | 37.25594, -6.99900 | SEV | 12 | 18 | 2 |

| | | | | | | | |
|---|---|---|---|---|---|---|---|
| 7 ES-DEHE | Spain. Seville, La Puebla del Río. Reserva Natural Concertada Dehesa de Abajo. | Shoreline of seasonal lake. | 37.20494, -6.17650 | SEV | 22 | 62 | 2 |
| 8 ES-PALM | Spain. Cadiz, Vejer de la Frontera. El Palmar de Vejer. | Coastal sand dunes. | 36.21777, -6.06055 | SEV | 2 | 88 | 2 |
| ES-GERE | Spain. Seville, Gerena. Arroyo de las Torres. | Edge of river bank. | 37.56250, -6.16000 | SEV | 0 | 13 | 0 |
| 9 UK-LEIG | United Kingdom. England, Leigh. | Edge of footpath over old mining waste. | 53.50191, -2.48386 | SEV | 6 | 0 | 0 |
| 10 UK-PENN | United Kingdom. England, Leigh. Pennington Flash. | Above the water line of the lake formed by coal mining subsidence. | 53.48277, -2.54864 | SEV | 5 | 7 | 1 |
| 11 UK-AIDA | United Kingdom. England, Leeds. RSPB St. Aidan's Nature Reserve. | Shoreline of lake formed from mining waste. | 53.74416, -1.41199 | SEV | 3 | 0 | 2 |
| 12 UK-FAIR | United Kingdom. England, Castleford. RSPB Fairburn Ings Nature Reserve. | Shoreline of lake formed from mining waste. | 53.74044, -1.32074 | SEV | 6 | 0 | 1 |

| | | | | | | | |
|---|---|---|---|---|---|---|---|
| 13 UK-MARS | United Kingdom. England, Southport. RSPB Marshside Nature Reserve. | Coastal marsh. | 53.67420, -2.97641 | SEV | 19 | 8 | 2 |
| 14 UK-MEER | United Kingdom. England, Ormskirk. WWT Martin Mere. | Freshwater lake. | 53.62133, -2.86725 | SEV | 19 | 5 | 1 |
| 15 UK-HOYL | United Kingdom. England, Wirral. Hoylake. | Edge of tidal marsh in Dee Estuary. | 53.39336, -3.18603 | SEV | 25 | 20 | 1 |
| 16 UK-DEVO | United Kingdom. England, Exminster. RSPB Exminster Marshes. | Damp muddy ditch in field. | 50.66467, -3.47143 | SEV | 7 | 0 | 1 |
| 17 IT-SCLY | Italy. Sicily. Palermo, Ficuzza. | Path in open woodland used for cow or cattle pasture. | 37.88523, 13.37004 | WHB | 31 | 0 | 1 |
| 18 IT-MADO | Italy, Sicily. Palermo, Parco delle Madonie. | Roadside. | 37.87827, 14.07768 | WHB | 15 | 0 | 0 |
| 19 IT-CAST | Italy, Sicily. Palermo, Castelbueno. | Roadside. | 37.91937, 14.11700 | WHB | 11 | 0 | 0 |
| 20 IT-POLL | Italy, Sicily. Palermo, Fiume Pollina. | In humid depressions and along path in the wide dry river bed. | 38.00537, 14.17702 | WHB | 21 | 0 | 0 |

| | | | | | | | |
|---|---|---|---|---|---|---|---|
| 21 NL-AMST | The Netherlands. Amsterdam, De Hoge Dijk. | Protected area. | 52.28691, 4.96577 | SEV | 17 | 10 | 1 |
| 22 NL-LEEU | The Netherlands. Leeuwarden, De Kleine Wielen. | Urban park. | 53.21470, 5.86810 | SEV | 10 | 17 | 1 |
| 23 AT-APET | Austria. Burgenland, Badesee Apetlon. | Lakeshore. | 47.78387, 16.83710 | WHB | 5 | 0 | 0 |
| 24 AT-AGGS | Austria. Niederösterreich, Aggsbach Markt. | Mud bank in a small bay on the Danube. | 48.27610, 15.38518 | WHB | 5 | 0 | 0 |
| 25 AT-RÜHR | Austria. Niederösterreich, Rührsdorf. | Muddy embankment to the Danube tributary. | 48.40303, 15.48838 | WHB | 11 | 0 | 0 |
| 26 AT-WALL | Austria. Niederösterreich, Wallsee. | Mudbank at the end of the oxbow. | 48.15948, 14.67543 | WHB | 19 | 0 | 0 |
| 27 CZ-KARD | Czechia. Kardašova Řečice, Velký řečický rybník. | Sandy banks of fishpond without water. | 49.18252, 14.86399 | WHB | 32 | 0 | 0 |
| 28 CZ-HAKL | Czechia. Haklovy Dvory, Novohaklovský rybník. | Drained marginal part of the fishpond with riparian vegetation and a lane. | 48.99301, 14.39617 | WHB | 6 | 0 | 0 |

| | | | | | | | |
|---|---|---|---|---|---|---|---|
| 29 DE-WIND | Germany. Schleswig-Holstein, Winderatter See. | Lakeshore. | 54.73878, 9.61435 | WHB | 5 | 0 | 0 |
| 30 DE-SÖRU | Germany. Schleswig-Holstein, Sörup-Dingholz. | Wet depression within an arable field free from crop. | 54.73934, 9.70182 | WHB | 25 | 0 | 0 |

| Table S2. Herbarium specimens identified as *Juncus minutulus* (Albert & Jahand.) Prain used in the morphological study. | |
|---|---|
| **Herbarium voucher** | **Location** |
| SEV5926 | Guadarrama, Spain |
| SEV10773 | Aldeanueva de Atienza, Spain |
| SEV42783 | Lac de Nino, Corsica, France |
| SEV150867 | Alosno, Spain |
| SEV150922 | |
| SEV150895 | Paymogo, Spain |
| SEV150897 | |
| SEV150898 | |
| SEV150907 | Valverde del Camino, Spain |
| SEV150916 | Zalamea la Real, Spain |
| SEV285937 | Fuenteovejuna, Spain |
| SEV285940 | Almodóvar del Río, Spain |
| SEV285941 | Arroyo Río Nevado, Spain |

| **Supplementary Table S3.** Average genome size and number of individuals (*n*) per ploidy level in populations sampled for flow cytometry. | | | |
|---|---|---|---|
| **Population code** | **Ploidy level** | **DNA content (pg/2C) (mean ± SD)** | ***n*** |
| ES-MART | 2x | 0.65 ± 0.09 | 4 |
| | 4x | 1.29 ± 0.11 | 2 |
| | 6x | 1.58 ± 0.06 | 10 |
| ES-DULC | 2x | 0.60 ± 0.04 | 9 |
| | 4x | 1.10 ± 0.00 | 1 |
| | 6x | 1.66 ± 0.07 | 4 |
| ES-DHER | 2x | 0.68 ± 0.01 | 9 |
| | 6x | 1.81 ± 0.06 | 11 |
| ES-RICE | 2x | 0.65 ± 0.04 | 75 |
| ES-ODIE | 2x | 0.62 ± 0.05 | 9 |
| | 6x | 1.65 ± 0.04 | 3 |
| ES-DEHE | 2x | 0.64 ± 0.04 | 12 |
| | 6x | 1.7 ± 0.11 | 10 |
| ES-PALM | 2x | 0.74 ± 0.00 | 1 |
| | 6x | 1.96 ± 0.00 | 1 |
| UK-LEIG | 6x | 1.69 ± 0.12 | 6 |
| UK-PENN | 6x | 1.71 ± 0.05 | 5 |
| UK-AIDA | 2x | 0.63 ± 0.08 | 2 |
| | 6x | 1.52 ± 0.00 | 1 |
| UK-FAIR | 6x | 1.75 ± 0.06 | 6 |
| UK-MARS | 2x | 0.64 ± 0.02 | 7 |
| | 6x | 1.76 ± 0.14 | 12 |
| UK-MART | 2x | 0.66 ± 0.00 | 1 |

| | | | |
|---|---|---|---|
| | 6x | 1.78 ± 0.04 | 18 |
| UK-HOYL | 2x | 0.67 ± 0.02 | 25 |
| UK-DEVO | 6x | 1.77 ± 0.07 | 7 |
| IT-SCLY | 4x | 1.08 ± 0.03 | 29 |
| | 6x | 1.55 ± 0.03 | 2 |
| IT-MADO | 4x | 1.14 ± 0.09 | 6 |
| | 6x | 1.77 ± 0.15 | 9 |
| IT-CAST | 2x | 0.59 ± 0.03 | 10 |
| | 6x | 1.64 ± 0.00 | 1 |
| IT-POLL | 2x | 0.61 ± 0.03 | 4 |
| | 6x | 1.62 ± 0.05 | 17 |
| NL-AMST | 6x | 1.73 ± 0.11 | 17 |
| NL-LEEU | 6x | 1.63 ± 0.08 | 10 |
| AT-APET | 2x | 0.59 ± 0.02 | 4 |
| | 6x | 1.68 ± 0.00 | 1 |
| AT-AGGS | 6x | 1.65 ± 0.03 | 5 |
| AT-RÜHR | 4x | 1.09 ± 0.01 | 4 |
| | 6x | 1.74 ± 0.12 | 7 |
| AT-WALL | 6x | 1.64 ± 0.04 | 19 |
| CZ-KARD | 6x | 1.67 ± 0.04 | 32 |
| CZ-HAKL | 6x | 1.68 ± 0.02 | 6 |
| DE-WIND | 6x | 1.97 ± 0.10 | 5 |
| DE-SÖRU | 6x | 1.83 ± 0.06 | 25 |

| **Table S4**. Putative morphospecies identification, DNA content and ploidy level of the ingroup individuals of the phylogenomic analyses. | | | |
|---|---|---|---|
| **Individual** | **Putative morphospecies** | **DNA content (pg/2C)** | **Ploidy level** |
| ES-MART1 | *Juncus hybridus* | 0.61 | 2 |
| NL-AMST2 | *Juncus bufonius s.str.* | 1.85 | 6 |
| NL-LEEU3 | *Juncus bufonius s.str.* | 1.82 | 6 |
| ES-MART4 | *Juncus bufonius s.str.* | 1.50 | 6 |
| ES-RICE5 | *Juncus hybridus* | 0.63 | 2 |
| ES-RICE6 | *Juncus hybridus* | 0.68 | 2 |
| UK-FAIR7 | *Juncus bufonius s.str.* | 1.77 | 6 |
| UK-AIDA8 | *Juncus bufonius s.str.* | 1.52 | 6 |
| UK-AIDA9 | *Juncus ranarius* | 0.68 | 2 |
| ES-DHER10 | *Juncus hybridus* | 0.68 | 2 |
| ES-DHER11 | *Juncus bufonius s.str.* | 1.85 | 6 |
| ES-DULC12 | Uncertain | 1.72 | 6 |
| ES-ODIE13 | *Juncus bufonius s.str.* | 1.66 | 6 |
| ES-ODIE14 | *Juncus ranarius* | 0.66 | 2 |
| IT-SCLY15 | *Juncus minutulus* | 1.11 | 4 |
| ES-DEHE16 | *Juncus hybridus* | 0.70 | 2 |
| ES-DEHE17 | *Juncus bufonius s.str.* | 1.69 | 6 |
| ES-RICE18 | *Juncus hybridus* | 0.59 | 2 |
| ES-RICE19 | *Juncus hybridus* | 0.70 | 2 |
| ES-RICE20 | *Juncus ranarius* | 0.69 | 2 |
| ES-RICE21 | *Juncus ranarius* | 0.62 | 2 |
| ES-MART22 | *Juncus minutulus* | 1.21 | 4 |
| ES-DULC23 | *Juncus ranarius* | 0.56 | 2 |
| ES-PALM24 | *Juncus bufonius s.str.* | 1.96 | 6 |
| ES-PALM25 | *Juncus hybridus* | 0.74 | 2 |
| UK-DEVO26 | *Juncus bufonius s.str.* | 1.83 | 6 |
| ES-DULC27 | *Juncus ranarius* | 0.64 | 2 |

| | | | |
|---|---|---|---|
| ES-RICE28 | *Juncus hybridus* | 0.58 | 2 |
| ES-RICE29 | *Juncus hybridus* | 0.59 | 2 |
| ES-MART30 | *Juncus bufonius s.str.* | 0.79 | 6 |
| UK-MEER31 | *Juncus bufonius s.str.* | 1.77 | 6 |
| UK-HOYL32 | *Juncus ranarius* | 0.66 | 2 |
| UK-MARS33 | Uncertain | 0.66 | 2 |
| UK-MARS34 | *Juncus bufonius s.str.* | 1.69 | 6 |
| UK-PENN35 | *Juncus bufonius s.str.* | 1.76 | 6 |

| **Table S5**. Outgroup sequence information | | | |
|---|---|---|---|
| **Family** | **Species** | **Material** | **Sample accession** |
| *Juncaceae* | *Juncus gerardii* | Silica dried | SAMEA6863453 |
| | *Marsippospermum gracile* | Herbarium | SAMEA112756269 |
| *Cyperaceae* | *Carex multispiculata* | Silica dried | SAMEA7754293 |
| | *Hypolytrum nemorum* | DNA bank | SAMEA7754195 |

**Table S6.** Results from Kruskal-Wallis tests.

| Variable | Chi-squared | df | p-value |
|---|---|---|---|
| CL | 4.2887 | 2 | 0.1171 |
| OTL | 2.1057 | 2 | 0.3489 |
| ITL | 7.7638 | 2 | 0.02061 |
| ROTC | 12.903 | 2 | 0.001578 |
| RITC | 29.064 | 2 | 4.89E-07 |
| ROTIT | 3.0644 | 2 | 0.2161 |

| Table S8.A. Pairwise comparisons of inner tepal length between ploidy levels | | | |
|---|---|---|---|
| **Comparison** | **Z** | **P-value (unadjust.)** | **P-value (adjusted)** |
| Diploid-Tetraploid | -1.3812622 | 0.167198358 | 0.50159507 |
| Diploid-Hexaploid | -2.6867194 | 0.007215755 | 0.02164726 |
| Tetraploid-Hexaploid | 0.3564599 | 0.721496144 | 1 |

| Table S8.B. Pairwise comparisons of ratio between capsule and outer tepal length between ploidy levels | | | |
|---|---|---|---|
| **Comparison** | **Z** | **P-value (unadjust.)** | **P-value (adjusted)** |
| Diploid-Tetraploid | -1.4822298 | 0.1382791498 | 0.414837449 |
| Diploid-Hexaploid | -3.5372919 | 0.0004042526 | 0.001212758 |
| Tetraploid-Hexaploid | 0.1255569 | 0.9000827026 | 1 |

| Table S8.C. Pairwise comparisons of ratio between capsule and inner tepal length between ploidy levels | | | |
|---|---|---|---|
| **Comparison** | **Z** | **P-value (unadjust.)** | **P-value (adjusted)** |
| Diploid-Tetraploid | -2.07401059 | 3.81E-02 | 1.14E-01 |
| Diploid-Hexaploid | -5.33490418 | 9.56E-08 | 2.87E-07 |
| Tetraploid-Hexaploid | 0.02432205 | 9.81E-01 | 1 |

**Supplementary Table S8.** Number of individuals with each seed ornamentation type per putative morphospecies.

| | **Smooth** | **Finely striated** | **Striated** | **Mixed** | **Unknown** |
|---|---|---|---|---|---|
| ***Juncus bufonius s.str.*** | 34 | 35 | 20 | 23 | 49 |
| ***Juncus minutulus*** | 4 | - | - | - | 62 |
| ***Juncus ranarius*** | 101 | 21 | 10 | 22 | 15 |
| ***Juncus hybridus*** | 49 | 10 | 2 | 12 | 6 |
| **Uncertain** | 21 | 15 | 2 | 13 | 20 |

| **Table S9.** Number of individuals per their disposition of flowers in the inflorescence | | |
|---|---|---|
| | **Solitary** | **Clustered** |
| ***Juncus bufonius s.str*** | 153 | 8 |
| ***Juncus minutulus*** | 60 | 5 |
| ***Juncus ranarius*** | 73 | 94 |
| ***Juncus hybridus*** | 3 | 76 |
| **Uncertain** | 47 | 24 |
